# The natural diversity of *E. coli* transporter-dependent capsules

**DOI:** 10.1101/2025.08.07.669119

**Authors:** Carine Roese Mores, Samuel Miravet-Verde, Elisabetta Cacace, Christoph Rutschmann, Chia-Wei Lin, Hans J. Ruscheweyh, Aline Cuenod, Elisa Cappio Barazzone, Enora Marrec, Kateryna Vershynina, Raffael Schumann, Dan J. Bower, Mario Schubert, Adrian Egli, Timm Fiebig, Emma Slack, Shinichi Sunagawa, Timothy G. Keys

## Abstract

Serotyping of bacteria using genomic information (*in silico* serotyping) has increasingly replaced serology. However, the *E. coli* capsule serotyping system has been largely abandoned since the 1990s, leaving gaps in our knowledge of capsule genetics, diversity, distribution, and epidemiology. To address this, we established a definitive genotype-serotype map for 35 serologically identified and structurally characterized transporter-dependent capsules. We then surveyed >37,000 *E. coli* genomes, cataloging 85 transporter-dependent capsule types (K-types), including 55 novel ones. We leveraged this catalog to develop an *in silico* serotyping tool, kTYPr, and applied it to curated sets of >25,000 *E. coli* genomes and metagenome-assembled genomes spanning diverse environmental and clinical sources. We found novel K-types enriched in under-sampled environments and associated with *E. coli* disease. This research expands our understanding of *E. coli* surface structures, supporting efforts for precision targeting with phage therapy or vaccines.

## Introduction

Antigens exposed on the bacterial surface are vulnerable to recognition by phages and host immune factors, driving extensive structural variation across bacterial populations. While the repertoire of glycans making up the O antigen of lipopolysaccharide, as well as large surface proteins such as adhesins and flagellin are well-studied,^1,2^ we have little information on the repertoire of capsular polysaccharides produced by *E. coli.*^3^ Capsular polysaccharides are considered virulence factors, owing to their ability to inhibit antibody binding to all other surface structures (including O antigens and proteins), as well as to block the action of innate immune mechanisms, e.g. opsonophagocytosis^4–6^ As the dominant exposed structure on the *E. coli* surface, capsular polysaccharides are therefore critical binding targets for protective antibodies and therapeutic bacteriophage, or even potential drug-targets.^5,7^

The constituents, linkages, and chemical modifications that make up capsular polysaccharides are highly diverse. During the last century, classical serology was used to identify 80 different capsule serotypes in *E. coli*.^8^ However, capsular polysaccharides are only weakly immunogenic, likely due to weak association to the bacterial surface.^9,10^ Correspondingly, antisera generated by vaccination with whole bacteria are dominated by other antibody specificities that cannot be completely removed by cross-absorption and agglutination is non-informative.^8^ Instead, counter-current immunoelectrophoresis (a precipitin-line-based approach) is required for capsule detection, which is slow, labor-intensive and expensive.^11^ With the advent of PCR-based *E. coli* typing of O- and H-antigens, research into capsule serology stalled, and the most recent new capsule serotype was described in 1977.^12^

*E. coli’s* capsular polysaccharides can be divided into four groups, based on their biosynthesis machinery and genetics.^13^ Group 1 and 4 capsules are known as Wzx/Wzy*-*dependent polysaccharides, these are synthesized in a manner analogous to O antigens. In contrast, group 2 and 3 capsules are ABC-transporter-dependent. These latter capsules are polymerized onto a phosphatidylglycerol oligo-β-3-deoxy-d-*manno*-octulosonic acid (Kdo) structure in the inner-leaflet of the inner-membrane and exported intact to the cell surface.^14–17^ The proteins required for oligo-β-Kdo synthesis (encoded by *kpsC* and *kpsS*)^14,16,17^ and capsule export (encoded by *kpsE*, *kpsD, kpsM,* and, *kpsT*) are well-conserved (Fig. 1A).^18^ K-type-specific genes extend the conserved oligo-β-Kdo, producing the serotype-specific part of the K antigen. Within this context, a dogma established in the field, based largely on knowledge of a few specific capsule serotypes such as K1 (associated with neonatal meningitis), states that group 2 and 3 capsules dominate extraintestinal pathovars, while group 1 and 4 capsules dominate in intestinal pathovars of *E. coli.* However, the lack of appropriate experimental and computational tools to identify capsule serotypes have hindered a comprehensive evaluation of these associations.^19^

**Figure 1.**
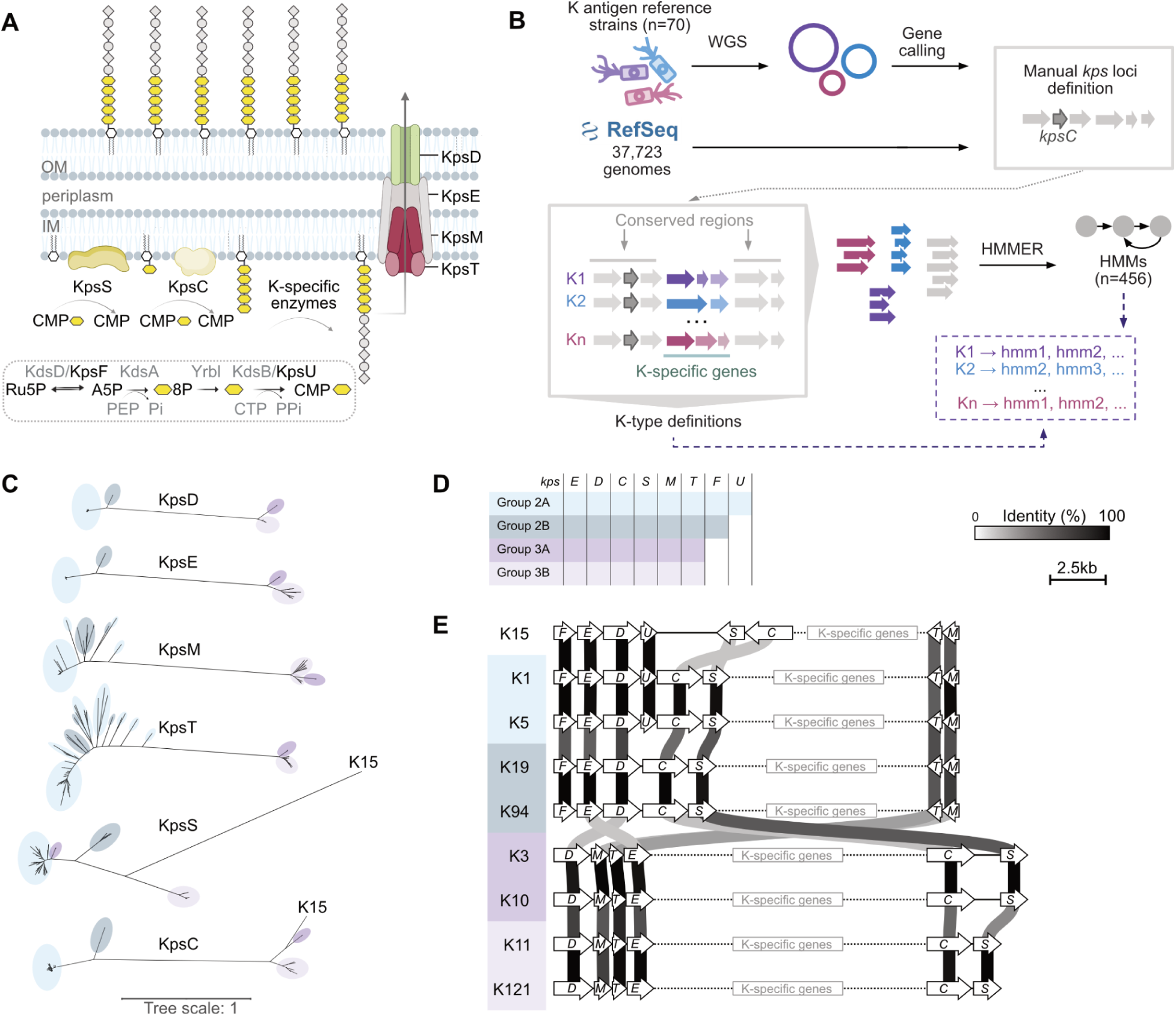
Overview of capsule biosynthesis, cataloging procedure, and *kps* lineages. (A) Schematic representation of transporter-dependent capsule biosynthesis and export, highlighting the roles of the conserved Kps proteins. (B) Construction of a comprehensive capsule gene cluster catalog by integrating K antigen reference strains (n=35) and RefSeq (n=37,723). The workflow includes whole genome sequencing (WGS) of reference strains, locus extraction, ORF identification, sequence clustering, and K-specific HMM generation (**STAR Methods**). (C) Unrooted phylogenetic trees for the essential proteins KpsEDCSMT involved in capsule biogenesis (STAR Methods). Leaves are colored according to the assigned lineage 2A, 2B, 3A, and 3B as in (D). (D) Presence/absence of conserved capsule biosynthesis genes in the four *kps* lineages. Colored bars indicate the presence of the corresponding gene. (E) Arrangement of conserved genes that is characteristic of each lineage. Two exemplary gene clusters from each lineage and the exceptional K15 cluster are shown. White arrows representing the conserved genes are drawn to scale. K-specific genes are indicated within each cluster (not to scale). Vertical greyscale lines indicate the percentage identity between the respective protein products.

With the increasing accessibility of whole-genome sequencing technologies and the escalating antibiotics resistance crisis, there is an urgent need for reliable methods to identify capsular polysaccharides from genetic data. Diverse computational approaches for *E. coli* serotype mapping from genomic data have been proposed, relying either on BLAST-based search^20–22^ or on more sensitive and robust Hidden Markov Models (HMMs).^23^ However, the absence of a comprehensive genotype-phenotype map limited the number of loci that could be linked with established capsule serotypes and prevented the assignment of novel K-types,^24–27^ leaving a critical knowledge gap.

Here we report major progress in closing this gap for the transporter-dependent class of capsular polysaccharides. We sequenced the *E. coli* capsule reference strain collection,^8^ confirming the transporter-dependent nature and establishing the genotype-to-serotype map for 35 known group 2 and 3 K antigens. We complemented this by cataloguing the diversity of capsule biosynthesis loci from 37,723 *E. coli* genomes, identifying 85 distinct gene clusters, including 55 putative novel transporter-dependent capsule types. We functionally annotated capsule biosynthesis genes by predicting protein structural similarities, allowing polysaccharide structural groups to be inferred for many novel K-types. For K-types with published polysaccharide structures, predicted gene functions aligned well with the expected biosynthetic requirements. However, we suggest revision of the K6 structure based on gene content and confirmative mass spectrometric analysis. The newly established catalog of genetically defined K-types was used to develop an HMM-based capsule typing tool, kTYPr. We demonstrated its utility by analyzing a globally distributed, non-redundant collection of 24,031 genomes and 2,763 stool-derived metagenome assembled genomes (MAGs). Our findings revealed a substantial overlap in K-type distribution between gut-colonizing and invasive *E. coli* strains, and uncovered associations between novel K-types and both human disease and understudied environments and hosts. We envision that the updated genotype-serotype reference and bioinformatic tool presented here will enable epidemiologists, infectiologists, vaccinologists and microbiologists to explore the role of *E. coli* capsules in infection, transmission and protection from disease.

## Results

### A genotype-serotype map for established capsule types

To establish a definitive K antigen genotype-serotype map we recently sequenced the reference strain collection established at the World Health Organization’s Collaborative Centre for Research and Reference on Escherichia.^8^ (Roese Mores et al., Genome Announcements, 2025) Genetic information for four further transporter-dependent K serotypes was obtained from publicly available sequence databases (K15 and K74) or from strains from the Culture Collection at the University of Gothenburg (K2a and K22). We identified a *kps* locus, including all essential genes (*kpsEDCSMT,* Fig. 1A), in 35 of the 70 strains, confirming their designation as ABC-transporter-dependent capsule types (groups 2 and 3).^3,13,18,28^ In contrast, these genes were not identified in strains designated as producers of Wzx/Wzy-dependent capsule types (groups 1 and 4).

In some reference strains we observed identical or near-identical *kps* loci (Fig. S1). These correspond to cross-reactive K serotypes,^8^ that consist of identical polysaccharide backbones with variable non-stoichiometric modifications.^29–34^ For example, K2ab is an acetylated version of the K2a backbone^32^. Accordingly, K2a and K2ab coding sequences differed by a single base insertion disrupting a putative acetyltransferase gene in K2a (Fig. S1A). Similarly, two missense mutations differentiate the candidate acetyltransferase genes in K13 and K23 (Fig. S1B), potentially reflecting differing acetylation of the respective polysaccharides.^35^ In some cases, the genes relevant to serotype differentiation are likely to be found outside of the capsule gene cluster (Fig. S1C-D).^36^ We have taken a cautious approach to assigning these closely related K-types where a robust genetic differentiation is not yet possible, grouping them under shared designations as follows: K2a and K2ab are assigned K2; K13 and K23 are assigned K13_K23; K18a, K18ab, and K22 are assigned K18_K22; K54 and K96 are assigned K96. Accordingly, we define a total of 30 K-types corresponding to the 35 reference transporter-dependent serotypes.

### An extended catalog of transporter-dependent capsules

Although previous studies have shown that there are more than the 35 established *kps* loci in *E. coli* genomes,^7,21,36,37^ the lack of a complete genotype-serotype map has prevented the clear identification of novel K-types. With this map now established, we set out to systematically characterize the diversity of transporter-dependent kps loci in *E. coli*.

To this end, we cataloged them in 37,723 *E. coli* genomes from the National Center for Biotechnology Information Reference Sequence Database (NCBI RefSeq; Fig. 1B, STAR Methods) by first using *kpsC* as a marker gene due to its role in biosynthesis of the unique oligo- β-Kdo that is characteristic of the glycolipid anchor common to this class of capsules.^14,16,17^ We then dereplicated sequences 30 kb up- and down-stream of *kpsC*, filtered for the presence of essential genes (*kpsEDCSMT*), and progressively trimmed the sequences to a presumed locus based on the organization of conserved genes, functional annotations, and identification of common flanking genes. The set of unique *kps* locus sequences was further refined using a protein catalog (excluding transposase-related ORFs), with each set of closely related ORFs (≥90% for both identity and coverage) used to build a protein HMM. In an iterative process, these HMMs were used to identify and extract the *kps* locus from *kpsC* flanking sequences (±30 kb), providing additional sequence diversity to build HMMs. A final set of 85 distinct *kps* loci was defined based on unique gene content, making our HMM-based K-type definitions robust to genetic rearrangements, and insertions or deletions in non-coding sequences. Comparison of the catalog with the reference strains led us to assign 55 putative novel K-types, named between K104 and K163, resulting in an updated catalog of 85 transporter-dependent K-types (STAR Methods).

### Defining a fourth lineage of transporter-dependent capsules

The historically designated group 2 and 3 capsules represent two distinct genetic lineages.^38^ A third group of plasmid-borne *kps* clusters (designated as 3B) was recently identified and proposed as a recent acquisition in *E. coli.*^7,36^ To investigate the evolutionary relationships across the 85 transporter-dependent K-types in our *kps* catalog, we constructed phylogenetic trees based on the conserved capsule biosynthesis proteins encoded by the *kpsEDCSMT* genes (Fig. 1C). Trees for the two most conserved proteins, the outer membrane pore (KpsD) and the periplasmic adapter (KpsE), revealed four distinct lineages. In addition to the three known lineages, we identified a previously unrecognized group which we designated 2B. These four lineages are further differentiated by the presence of *kpsF* and *kpsU*, which encode enzymes involved in CMP-Kdo biosynthesis and are redundant in the *E. coli* genome (Fig. 1A, D), and by the organization of conserved genes (Fig. 1E). Phylogenetic trees of KpsC and KpsS share the KpsDE-architecture with two known exceptions; the *kpsS* genes in the 3A lineage appear to derive from the 2A lineage;^36^ furthermore, KpsC and KpsS from K15 form distinct branches that are incongruent with the other lineages, suggesting a possible origin outside *E. coli.*^39,40^ Phylogenetic trees of KpsM and KpsT revealed higher diversity and evidence of exchanges between the 2A and 2B lineages, consistent with frequent rearrangements of *kpsM*, *kpsT* and the K-specific genes in these lineages, as observed in K7, K51, K52, K104, K105, K106, and K108 (Fig. 2).

**Figure 2.**
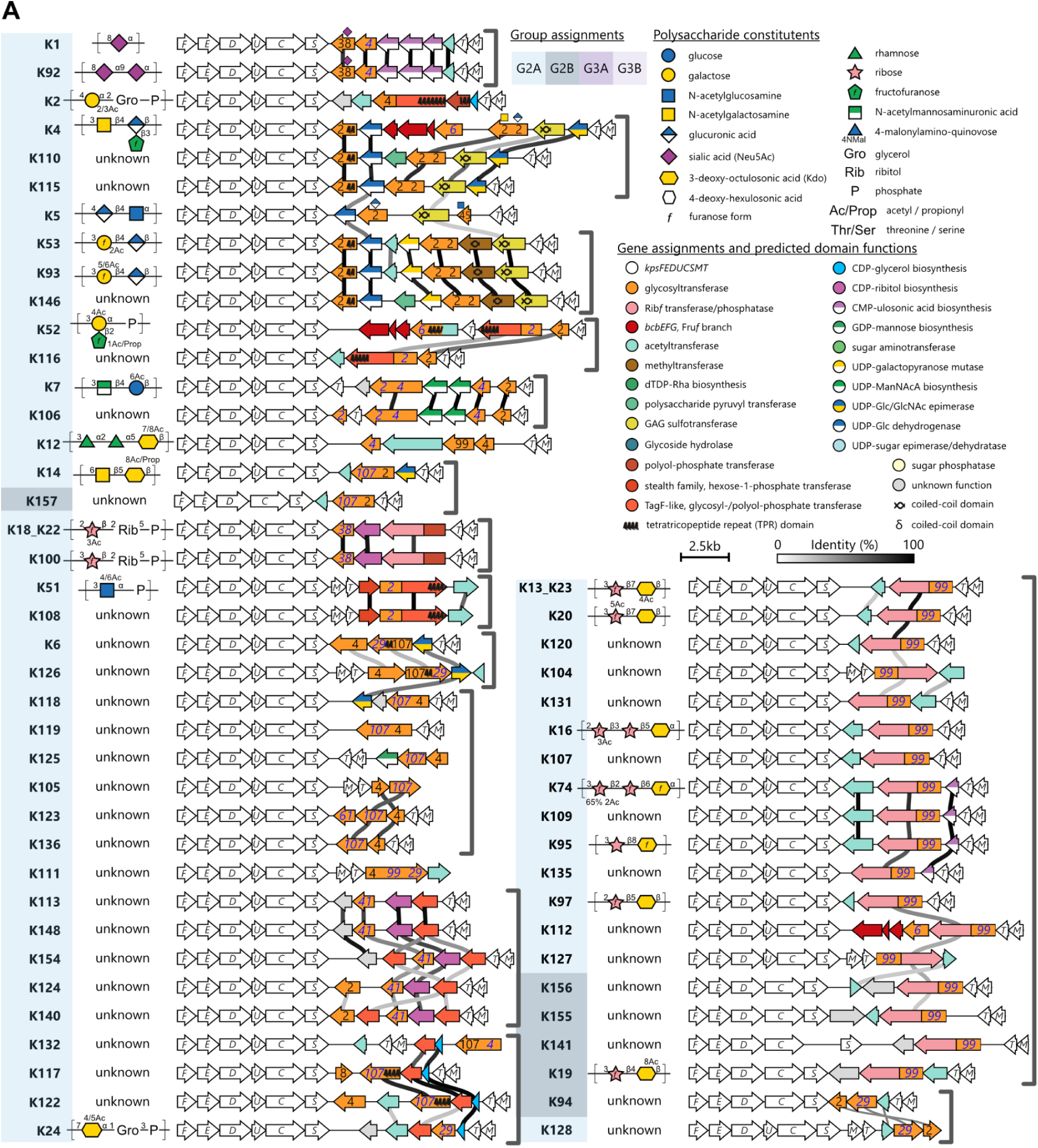
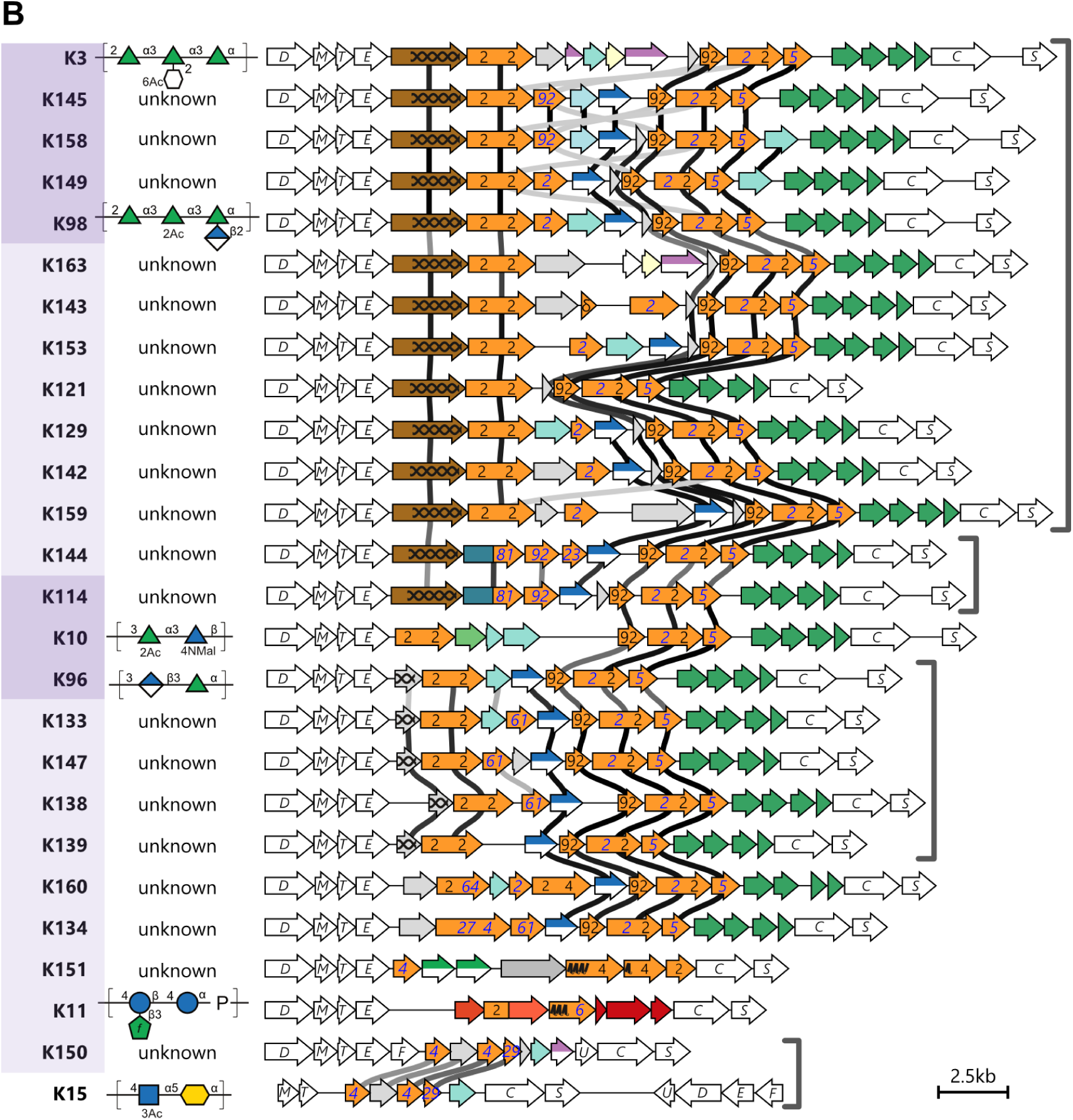
Catalog of *E. coli* transporter-dependent capsule gene clusters. The conserved biosynthetic and export genes are white. K-specific genes are colored according to known or predicted function. For characterized glycosyltransferases (GT), the transferred sugar is displayed above the gene. The CAZy family for GT domains are indicated in black numbers on the gene. For putative GT domains identified by predicted structural similarity, the CAZy family of the closest structural match is indicated in blue italics on the gene. Pairwise polypeptide identities >30% between vertically adjacent K-specific genes are indicated by grey-scale connecting lines. For clarity, sequence similarity between conserved genes is not shown. Gene clusters are organized to collect groups with functionally related gene content, indicated by grey brackets to the right of gene clusters. Polysaccharide repeat units, where known, are depicted according to SNFG conventions.^54^ (A) Gene clusters assigned to lineages 2A and 2B. (B) Gene clusters assigned to lineages 3A, 3B and K15.

### Functional annotation of K-specific genes is supported by predictions of protein structures

To explore the functional repertoire within the catalog, we functionally annotated serotype-specific genes. We identified conserved domains and glycosyltransferase (GT) families according to the Carbohydrate-Active enZYmes Database (CAZy, STAR Methods).^41–45^ We also predicted structural similarities with proteins in the Protein Data Bank (PDB), using AlphaFold2^46^ and Foldseek.^47^ Altogether these analyses assigned putative functions to 421 of 456 ORFs (Fig. 2). Sequence similarity allowed us to assign a CAZY family to 50.5% (137 of 271) of the GT domains. The remaining GTs were assigned based on predicted structural similarity with characterized CAZy family members in the PDB, providing functional insights in the absence of clear sequence similarity. For example, 29 gene clusters encode a domain with predicted structural similarity to the related GT99 and GT107 families,^14^ whose characterized members are retaining Kdo transferases (KdoT). This aligns with the presence of Kdo in the polysaccharide repeat units of clusters with known structures (namely, K13_K23, K14, K16, K19, K20, K74, K95 and K97), and suggests a role for GT99/107-like domains in catalyzing the respective Kdo linkage.

Grouping *kps* clusters with functionally related K-specific genes revealed 18 sets of K-types predicted to encode polysaccharides with related structural features, e.g. similar backbone composition (Fig. 2). Thirteen of these sets include at least one cluster with a known polysaccharide structure, facilitating structural inferences for novel polysaccharides. A striking example is the set of K-types (K13_K23 to K19, Fig. 2A) sharing a large ORF (>1100 amino acids) with little detectable sequence similarity but a consistent set of three domains identified by predicted structural similarity: a GT99-like KdoT, and a dual domain ribofuranosyltransferase/phosphatase, recently shown to be the biosynthetic origin of ribofuranose (Rib*f*) in bacterial polysaccharides.^48,49^ These domains are consistent with Kdo- and Rib*f*-containing polysaccharides published for seven members of the group. This example illustrates the power of predicted structural similarity (as opposed to sequence similarity) to support functional annotation of carbohydrate active enzymes by identifying functional relationships obscured by sequence divergence, and to provide structural hypotheses for novel capsular polysaccharides.

### Structural analysis of polysaccharides clarifies inconsistencies between functional gene content and published structures

Comparison of the predicted functions in each gene cluster with published polysaccharide structures revealed broadly consistent associations across the catalog (Fig. 2), with two notable exceptions. In the first case, the K6 gene cluster lacks predicted Rib*f*T domains but is reported to produce a Rib*f*-Rib*f*-Kdo repeat unit.^50^ To clarify this inconsistency, we purified capsular polysaccharide from seven reference strains reported to produce various Rib*f*-Kdo polysaccharides and analyzed repeat unit composition by MALDI-MS of permethylated glycans (STAR Methods, Fig. S2). Consistent with our expectations based on the gene clusters, we observed Rib*f*-Kdo repeat units in all except the K6 polysaccharide. There we observed m/z species consistent with a homopolymer of Kdo. Notably, a putative poly-Kdo structure is in agreement with the presence of an ORF encoding a dual GT29-GT107 domain protein, both of which are known for transfer of ulosonic acids, such as Kdo, from CMP-activated donor substrates. We account for these results by assigning K6 as a serotype with an unknown polysaccharide structure (Fig. 2) pending further structural investigation.

On the other hand, K16 is reported to produce a Rib*f*-Rib*f*-Kdo trisaccharide repeat unit^51^ despite having an identical set of predicted domains to several clusters reported to produce a Rib*f*-Kdo disaccharide repeat (e.g. K13, K19, K20, etc., Fig. 2). MALDI-MS analysis of permethylated glycans revealed the expected disaccharide repeats for K13, K19, K20, K23, and K97 and, contrary to our expectations, the Rib*f*-Rib*f*-Kdo trisaccharide repeat unit was observed for K16 (Fig. S2). The presence of this trisaccharide repeat in K16 indicates a gap in our biochemical understanding. It may be explained by either a bifunctional Rib*f*T capable of transferring Rib*f* to both Kdo and Rib*f* acceptors, or the involvement of an additional Rib*f*T encoded outside of the capsule gene cluster. Notably, the presence of single-domain bifunctional glycosyltransferases involved in biosynthesis of other capsules favors the former hypothesis. These include a GlcNAc transferase that transfers two consecutive GlcNAc moieties in forming the *Neisseria meningitidis* serogroup L capsule^52^ and the *E. coli* K92 polysialyltransferase which transfers successive sialic acid residues with alternating linkages.^53^

### Development of kTYPr as a genotype-to-serotype profiler in *E. coli*

We used the newly assembled *kps* catalog and HMMs to develop kTYPr, an *in silico* capsule typing software applicable to both genomic and metagenomic data (Fig. 3, STAR Methods). The tool screens genomes for *kpsC* gene presence, and processes *kps*-positive ones with the full set of HMMs to identify conserved and K-type specific genes that meet HMM-specific bit score cutoffs. K-types are defined by the presence of all essential genes (*kpsEDCSMT*) and K-type specific gene sets. In cases where criteria for multiple K-types are met, the decision is based on the highest accumulated bit scores. This allows the tool to distinguish between closely related gene clusters (e.g. K51 and K108) and to avoid incorrectly assigning K-types that are defined by a subset of genes that are also present in other K-types (e.g. K121 is a subset of K153 and K129, see Fig. 2). Bit score values are reported for all K-types considered, together with the complete set of hits for every gene against our HMM collection for further inspection or parametrization, a genbank file of the extracted *kps*-related genes, and a visualization of its similarity to the predicted or closest K-type cluster using clinker^55^ (Fig. 3A).

**Figure 3.**
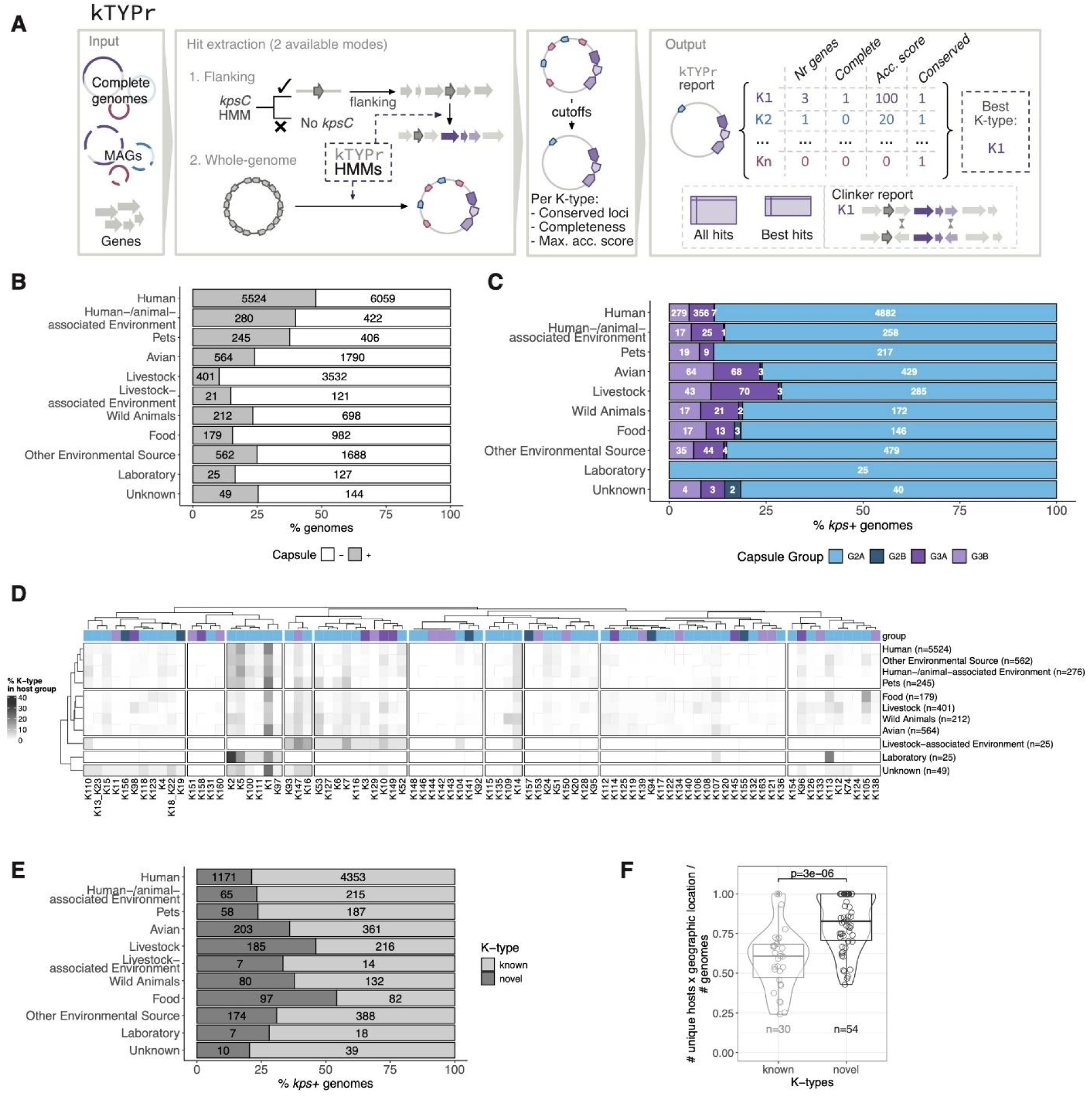
kTYPr workflow and K-type diversity across hosts and environments. (A) Schematic representation of the kTYPr tool, highlighting allowed inputs (complete genomes, MAGs or genes) and two hit extraction modes (flanking and whole-genome). Once the custom cutoffs are applied, the hits are processed to produce a report output including for each input genome the accumulated bit-scores and completeness metrics for all K-types profiled, with the “best match” as the most complete K-type with highest accumulated bit-score. In addition, all gene hits against the HMM collection, a formatted genbank and a clinker visualization of the similarity between the genome and the best hit(s) are also reported (STAR Methods). (B) Fraction of capsule*-*positive genomes (defined as containing a complete *kps* locus, STAR Methods) across hosts and environments. Absolute counts are indicated within the bars. (C) Capsule group proportion across hosts and environments. Only capsule-positive genomes are considered. Absolute counts are indicated within the bars. (D) K-type proportion in hosts and environments. Only capsule-positive genomes were considered. K-types and hosts are clustered according to Spearman correlation. K-type groups are color-coded as in Fig. 3C. (E) Proportion of novel and known K-types across hosts and environments. Only capsule-positive genomes are considered. (F) Number of distinct hosts and geographic locations (divided by number of genomes) for each K-type. The p-value between known and novel K-types is shown (two-sided Wilcoxon test); box limits correspond to first and third quartiles, with the median marked, and whiskers extending to the most extreme data points up to 1.5 times the interquartile range (IQR). Data point counts are indicated below the boxplots.

We evaluated the performance of kTYPr on the source dataset (37,723 *E. coli* genomes from RefSeq). Of these, 25.5% (n=9,607) were assigned a transporter-dependent K-type based on the presence of the complete set of essential genes (*kpsEDCSMT*) and a complete set of K-type-specific genes (Fig. S3A). Of the 28,082 genomes that were not assigned a transporter-dependent K-type using default settings (STAR Methods), 92.2% (n=25,879) lacked a complete set of essential and K-specific genes. The remaining 5.1% (n=1,436) had a complete set of either *kpsEDCSMT* or K-specific genes, but not both. These cases may represent disrupted gene clusters, genes missing in draft genomes, or uncataloged *kps* cluster diversity. While all 85 K-types were identified in the source dataset, 11 were only identified once or twice (Fig. S3B), indicating that our catalog has deeply sampled the diversity present in RefSeq, but that RefSeq has not sampled the diversity of *E. coli kps* loci present in nature.

### Novel K-types are more represented in understudied hosts and environments

To investigate the distribution of group 2 and 3 capsules across environments, hosts and body sites in humans, we used kTYPr to profile a collection of 32,043 metadata-curated, globally distributed genomes from NCBI (STAR Methods). To reduce genomic redundancy due to the isolation of clonal strains from the same sample, or repeated isolation of strains from the same individual over time, we dereplicated all genomes coupling a 99% ANI cutoff with metadata on sample origin (host type, sample type, geography, health state) and bacterial characteristics (sequence type, O-, H- and K-type), resulting in 23,206 representative genomes. We further integrated this dataset with a collection of 825 *E. coli* genomes from urinary tract infection (UTI) cases, 259 of which were associated with invasive disease (STAR Methods).^56^

The resulting collection (n=24,031) was biased towards human hosts (48.2%, Fig. 3B) and westernized geographical areas (49.7% of genomes from Europe and North America, Fig. S4A). 33.5% of the genomes (n=8,062) contained a complete *kps* locus, with a higher proportion in humans (47.7%), pets (37.6%) and human-associated environments (e.g. houses, public transportation, sewage water and hospitals, 39.9%) (Fig. 3B). Most of the assigned transporter-dependent capsules belonged to group 2 (86% of genomes, n=6,958), of which only 25 genomes belonged to group 2B. 1,104 genomes (13.7%) harboured group 3 capsules, of which 609 genomes of group 3A and 495 of group 3B (Fig. 3C). Humans, human-associated environments and pets showed highly correlated K-type profiles (Fig. 3D, S4B-C), whereas livestock and wild animals were more similar, in agreement with previous reports of genetically distinct *E. coli* lineages between humans and livestock.^57^ Animal hosts that are well-represented in the collection correlate according their K-type profiles, beyond geographical groups (Fig. S4C).

While 78.7% of the K-types in highly studied human hosts and associated environments were previously reported, the new K-types identified in this study were more frequent in less studied environments, such as non-domesticated animals (Fig. 3D-E). Novel K-types were also more diverse in terms of hosts and geography than known ones (Fig. 3F). The most frequent K-types overall were K1, K5, K2 and K14 (Fig. S4D). This ranking reflected the bias of the collection towards human samples and, except for the generally most abundant K1 type, was not always conserved in other environmental and geographical niches (Fig. S4E): for example K105 and K109 (newly defined types) were first and second most frequent in food (13.4%) and wild animals (7.1%), respectively (Fig. S4B). Overall, our approach unveiled a much higher diversity of transporter-dependent capsules than previously known from highly sampled, human-associated environments.

### Diversity of transporter-dependent capsules in human health and disease

Consistent with previous reports,^58^ 78% of *E. coli* genomes from phylogroups B2, D, and F harboured group 2 K-types (Fig. S5A). This was largely driven by human samples (Fig. 4A), where these phylogroups have been associated with extraintestinal pathogenic *E. coli* (ExPEC) (Fig. S5B).^59^ Because most sequenced *E. coli* genomes are from clinical isolates, we have limited understanding of the prevalence and distribution of K-types in asymptomatic individuals. To address this gap, we additionally profiled a globally-distributed collection of *E. coli* metagenome-assembled genomes (MAGs) from the stool of healthy humans from 25 published studies (n=2,763 of which 921 *kps*-positive, Fig. S5C).^60^

**Figure 4.**
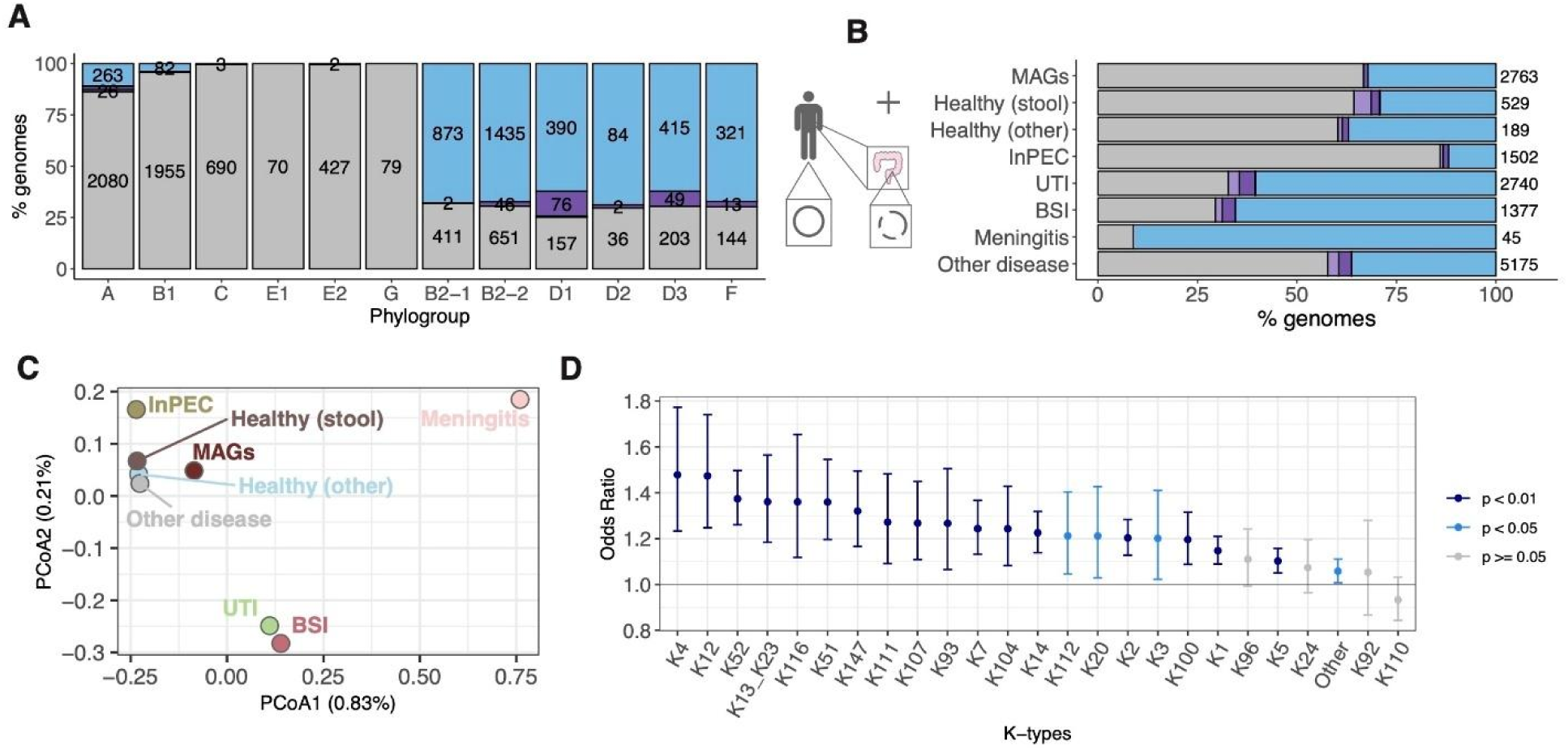
K-type diversity in human health and disease. (A) Proportion of capsule groups across phylogroups in *E. coli* complete genomes from humans (n=11,537). Groups are color-coded as in Fig. 3C, with grey indicating *kps*-negative genomes. (B) Proportion of capsule groups in *E. coli* genomes from humans across clinical categories and in asymptomatic carriers (MAGs from stool of healthy individuals). Groups are color-coded as in Fig. 3C. Non-*E. coli*-associated diseases are grouped as “other disease”. InPEC, Intestinal Pathogenic *E. coli*; UTI, Urinary Tract Infection; BSI, bloodstream infection. (C) Principal Coordinates Analysis (PCoA) of K-type profiles from the clinical categories in 4B. The PCoA was based on a Bray-Curtis dissimilarity matrix calculated from the proportion of K-types in each group. (D) Odds ratio (OR) and confidence interval (CI) from a generalized linear model evaluating the contribution of K-types to invasiveness. A dataset consisting of asymptomatic carriage (2,763 *E. coli* MAGs from healthy individuals, and 773 complete genomes from stool of healthy individuals) and invasive *E. coli*-associated disease (1377 genomes of which 1,181 genomes of isolates from blood or cerebrospinal fluid from NCBI and 259 from blood from a published study on urosepsis)^56^ was used (STAR Methods). OR=1 is indicated (dashed line). Data points are color-coded according to statistical significance (STAR Methods).

Although group 2 capsules were also more frequent in MAGs belonging to phylogroups B2, D and F (Fig. S5D), their proportion was higher in extraintestinal disease (e.g. isolates from blood or cerebrospinal fluid) (Fig. 4B, S5D). Phylogroups B2, D and F had also the highest K-type diversity, higher in invasive disease than in asymptomatic carriage, except for phylogroup B2-2 (Fig. S5E-G). K-type profiles were more similar among MAGs and genomes from healthy individuals, patients with diseases not associated with *E. coli* and patients with intestinal pathogenic *E. coli* (InPEC), while genomes from urinary tract infections (UTIs), bloodstream infections (BSIs) and meningitis showed distinct separation (Fig. 4C). Transporter-dependent capsules were less common in InPEC (14%) compared to extraintestinal disease (67.3% in UTIs, 70.5% in BSI, 91.1% for meningitis), and to stool-derived MAGs (33.3%) or genomes (35.7%) from healthy individuals (Fig. S5H). Altogether, while consistent with previous reports of group 2 K-types in ExPEC,^61,62^ our analysis revealed the extensive presence of typical ExPEC K-types (e.g. K1, K2, K5)^19,63,64^ in the gut microbiome of asymptomatic individuals by overcoming previous limitations of isolate-based reference databases.

Furthermore, in agreement with the low frequency of transporter-dependent capsules in InPECs (Fig. S5H), we did not detect transporter-dependent K-types in genomes associated with the typical InPEC serotypes O157:H7 (enterohemorrhagic *E. coli*, EHEC) and O104:H4 (enteroaggregative *E. coli*, EAEC) (Fig. S6A). We note that these strains may harbour group 1 or 4 capsules that are not detected by kTYPr. On the other hand, typical ExPEC serotypes, such as O25:H4 and O1:H7, had the highest number of distinct transporter-dependent K-types and were among the O-types with highest proportion of capsule positivity (Fig. S6A). O- and K antigen co-occurrence may also reveal functional interdependencies between glycan biosynthetic gene clusters: for example, K12 polysaccharide contains rhamnose.^65^ However, since we could not find rhamnose-biosynthesis genes in the *kps* cluster as for other rhamnose-containing K-types (Fig. 2), we hypothesized that these K-types might harness the rhamnose-biosynthesis machinery of the O antigen. Consistent with this hypothesis, K12 was exclusively found with rhamnose-containing O-types (O4, O139, O51, O1, O132, O13/O135, O18, O25) (Fig. S6B). Overall, this demonstrates how kTYPr can inform new hypotheses on functional relationships between different bacterial surface sugar antigens with potential relevance for human disease.

Finally, to quantify the association of K-types with invasiveness compared to asymptomatic carriage, we used a generalized linear model, with the invasive phenotype (defined as isolation from blood or cerebrospinal fluid in patients with bloodstream infection or meningitis) as binary outcome, K-type, O-type, H-type, sequence type (ST), phylogroup, host age, gender, and geographical location as fixed effects (STAR Methods). UTI cases were excluded to rule out possible contamination from non-pathogenic *E. coli* from the gut. The model confirmed previously proposed associations, including K1, K2, K4 and K5.^19,63,64^ The K-types with the strongest association with invasiveness were K4, K12, K52 and K13_K23. Of the 20 K-types that were significantly and positively associated with invasiveness, 10 were previously reported (of which five only recently).^37^ Of the 10 types that we reported for the first time with invasive potential, six were among those newly defined in our study: K104, K107, K111, K112, K116 and K147 (Fig. 4D), highlighting novel contributors to invasiveness that should be investigated in larger datasets and mechanistic studies.

## Discussion

Capsular polysaccharides have historically been classified through serology and bacteriophage sensitivity, but these approaches have inherent limitations in resolution, scalability and reproducibility.^8,11,66,67^ Our work advances the field by integrating genomic classification, linking defined sets of genes to both known and putative novel serotypes. Based on a ground-truth database including all established *E. coli* K antigens, and an extended catalog of *kps* clusters assembled from >37’000 *E. coli* genomes, we provide a high-throughput, scalable and robust genome-driven approach to studying *E. coli* transporter-dependent capsules.

Our study significantly expands the known diversity of *E. coli* transporter-dependent capsules. Our catalog of 85 K-types, including 55 putative novel types (Fig. 2) confirms the idea that *E. coli* capsule diversity extends far beyond the set collected by serologists in the 20th century.^7,19,21,36,37^ Phylogenetic analysis of the conserved capsule biosynthesis machinery clarifies the evolutionary relationships underlying the group 2 and 3 classifications (Fig. 1C-E). Our work also revealed a previously unrecognized lineage, group 2B, characterized by the absence of *kpsU*. We confirm and expand the group 3B lineage, first identified by Hong et al.^7,36^ as plasmid-borne *kps* clusters. While we have not investigated the presence of *kps* clusters on mobile genetic elements, our work provides a method to explore this in future studies.

A key contribution of this study is the comprehensive characterization of the *kps* cluster catalog, assigning putative functions to more than 90% of K-specific ORFs (Fig. 2). An earlier study of TagF-like domains in capsule polymerases showed that sequences with as little as 15% identity shared fold and function (sugar-phosphate and polyol-phosphate transfer).^68^ Building on this, we integrated structure prediction and similarity searches into our annotation pipeline, almost doubling the number of glycosyltransferase domains identified and associated with a GT family in the CAZy database. As our MALDI-MS results for K6 and K16 demonstrate (Fig. S2), this functional and structural map can be used to inform mechanistic hypotheses on capsule biosynthesis, to highlight gaps in our understanding of polysaccharide structure and biochemistry, and to infer structures for novel K-types.

To open this resource for the research and public health communities, we developed kTYPr, an HMM-based tool for robust and scalable determination of K-types from genome sequences. Unlike earlier BLAST-based approaches,^21,37^ kTYPr uses gene content-based K-type definitions and sensitive HMMs for more accurate detection. These advances allow kTYPr to recognize gene clusters that are highly divergent, reorganized, distributed across different genomic loci, or fragmented in MAGs. The critical resource that sets kTYPr apart from other available tools is our ground-truth dataset generated by sequencing the available reference strains. This makes kTYPr the first tool that accounts for all 35 known transporter-dependent K-types (corresponding to 30 distinct gene clusters) and differentiates 55 putative novel types.

A limitation is that kTYPr’s accuracy is inherently constrained by our understanding of the genetic and biochemical determinants of capsule structure. For example, K18 and K22 share identical capsule gene clusters, yet differ in acetylation of the polysaccharide (Fig. S1C). As the responsible acetyltransferase gene is not located within the capsule locus, and remains unidentified, we currently lack the knowledge to distinguish these serotypes using genomic data alone. Similar ambiguities arise in other cases, such as K2a/K2ab and K13/K23. These examples highlight the need to advance our understanding of capsule biosynthesis, and to develop methods that increase the speed and throughput of structural analysis of bacterial polysaccharides. Integration of these tools with genomic data will be essential to future advances in genome-driven prediction of bacterial surface structures.

This work illustrates the power of kTYPr, applied to a unique collection for global distribution, metadata resolution and genome dereplication, revealing type-resolved capsule ecology. We found transporter-dependent capsules across diverse hosts and environments (Fig. 3B and S4C), but the distribution of capsule lineages and types differs substantially between them (Fig. 3C-D). The 30 known types were on average less ecologically diverse than 55 novel ones, which were more often found in undersampled hosts, such as wild animals (Fig. 3E-F). For example, half of the K-types identified in livestock and food have not been previously described and the newly described K105 and K109 were the most frequently observed types after K1 in livestock and wild animals, respectively (Fig. 3D). Our initial screen on available strains with associated metadata hints at associations between transporter-dependent capsules and hosts and potentially geographical location (Fig. S4C, E), which should be further investigated with larger datasets representing undersampled hosts such as wild animals.

The inclusion of *E. coli* MAGs from healthy individuals as carriage control revealed previously unexplored associations of *E. coli* capsules with human health and disease. An important finding is the broadly similar distribution of K-types among *E. coli* isolates from the healthy gut and from clinically infected human patients (Fig. 4C, S5F). These results confirm, at unprecedented scale, the observations of serologists that ExPEC-associated serotypes are good colonizers of the intestinal tract.^69^ These observations also support recent analyses suggesting that opportunistic infection of vulnerable hosts might explain a large portion of ExPEC-associated K-type diversity.^37^

Overall, we presented here a scalable, fast and flexible tool to type transporter-dependent capsules, kTYPr, and introduced three new resources: a well-characterized catalog of known and novel *kps* loci based on a ground-truth genotype-serotype map for all 35 reference transporter-dependent capsule types, and two curated, globally distributed, and dereplicated collections of *E. coli* genomes and MAGs, which can be used to revisit our current knowledge of *E. coli* ecology and genetic diversity at unprecedented scale. Our results enable structural inference for novel capsule types, highlight key gaps in our knowledge of polysaccharide structure and biochemistry, and outline the diversity of capsules across ecological niches, including novel K-types that are associated with understudied environments and invasive disease. This study provides a foundation for future work on *E. coli* glycobiology, ecology, evolution, epidemiology and for the prevention of ExPEC infections with capsule-based vaccines,^58^ and treatment of infections with capsule-dependent monoclonal antibodies^70^ or phages.^71^

## Acknowledgments

This work was supported by funding from the Basel Research Centre for Child Health (BRCCH) to E.S. and S.S., the Swiss National Science Foundation through the project grants 40B2-0_180953 and 310030_185128 to E.S. and the NCCR Microbiomes (51NF40_180575 and 51NF40_225148) to E.S. and S.S, and an Innosuisse grant (117.143 IP-LS) to T.G.K and E.S., and core funding from ETH Zürich. E.S. acknowledges support of an European Research Council Consolidator Grant (865730). E.S. and S.S. are supported by the LOOP Zurich mTORUS project. S.M.V. acknowledges funding from the Human Frontier Science Program (HFSP) through a postdoctoral fellowship [LT0050/2023-L]. E.C. acknowledges support from SNSF Spark (CRSK-3_228959) and Novartis Freenovation (FN24-0000000703). AlphaFold runs were performed on the ETH Euler cluster. We acknowledge the support of the IT Service and HPC facilities of ETH Zürich.

Conceptualization (E.S., S.S., T.G.K.), Data curation (C.R.M., E.C.), Formal analysis (C.R.M., S.M.V., E.C., T.G.K.), Funding acquisition (E.S., S.S., T.G.K.), Investigation (C.R.M., S.M.V., E.C., C.R., K.V., C-W.L., E.C.B, E.M., R.S., T.F., M.S., T.G.K.), Methodology (C.R.M., S.M.V., E.C., D.B., T.G.K.), Software (D.J.B., S.M.V.), Resources (H.J.R., A.C., A.E.), Supervision (E.S., S.S., T.G.K.), Validation (S.M.V., E.C., T.G.K.), Visualization (C.R.M., S.M.V., E.C., T.G.K.), Writing – original draft (C.R.M., S.M.V., E.C., T.G.K.), Writing – review & editing (C.R.M., S.M.V., E.C., E.S., S.S., T.G.K.). A.E. was supported by an unrestricted research grant by the University of Zurich.

## Declaration of generative AI and AI-assisted technologies in the writing process

During the preparation of this work the authors used ChatGPT to improve readability of the manuscript. The authors reviewed and edited all content and take full responsibility for the content of the published manuscript.

## Supplementary Figures

**Figure S1.**
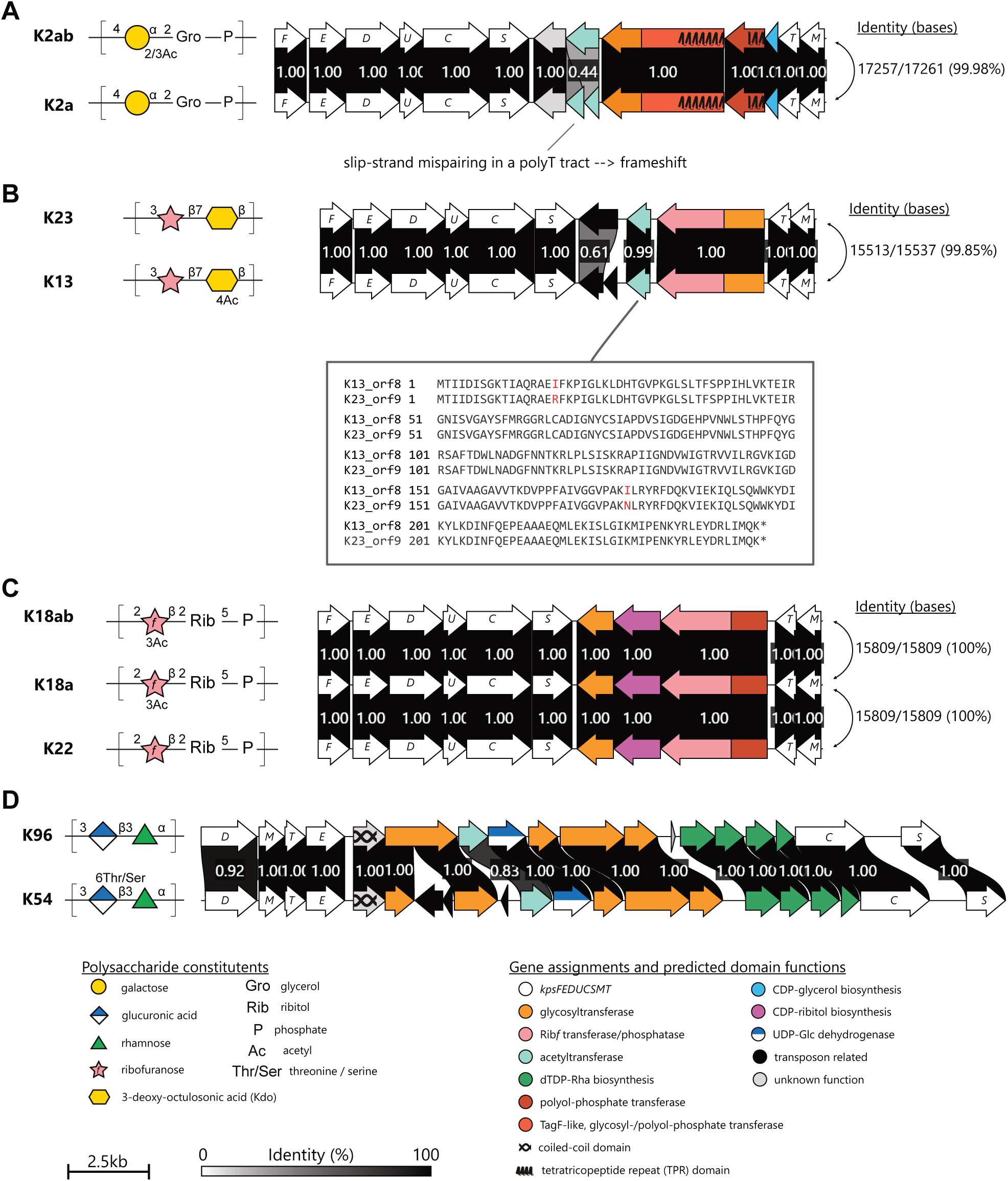
Closely related capsule gene clusters identified in different K serotype reference strains do not support robust genetic differentiation of the respective serotypes, related to Figure 2. Capsule gene clusters were compared and rendered using clinker. The proportion of identical amino acids is displayed numerically and in grey-scale between opposing ORFs. The reported base identities are from global Needle-Wunsch alignments and the protein alignments were made with the Smith-Waterman algorithm. Gene assignments and functional predictions were made as described in STAR Methods. Polysaccharides are depicted according to the conventions of the Symbol Nomenclature for Graphical Representation of Glycans and references for the displayed polysaccharide structures are provided in. (A) K2a and K2ab differ by acetylation of galactose in the repeat unit, consistent with a frameshift mutation in the putative acetyltransferase ORF in K2a. (B) The K13 and K23 polysaccharides differ by acetylation of the Kdo residue in the repeat unit and the candidate acetyltransferases differ by two missense mutations. Although the observed mutations may explain the K2a/K2ab and K13/K23 serotypes, they do not provide a robust basis for genetically distinguishing these serotypes. Therefore, these serotypes are grouped under the names K2 and K13_K23, respectively, in our catalog. (C) The K18 polysaccharide is an acetylated version of the K22 backbone, but the capsule gene clusters are identical (100% identity), and there is no candidate acetyltransferase gene within the clusters. The serological distinction between K18a and K18ab is unclear because chemical analysis of the polysaccharides revealed identical repeat unit structures. As there is currently no genetic basis for distinguishing these serotypes, they are grouped under the assignment K18_K22 in our catalog. (D) Our analysis of the K54 and K96 gene clusters is consistent with earlier reports suggesting that the gene(s) responsible for threonine modification of the polysaccharide backbone are located outside of the kps locus. Pending identification of these gene(s), the K54 and K96 clusters are grouped under the assignment K96 in the current version of the catalog.

**Figure S2.**
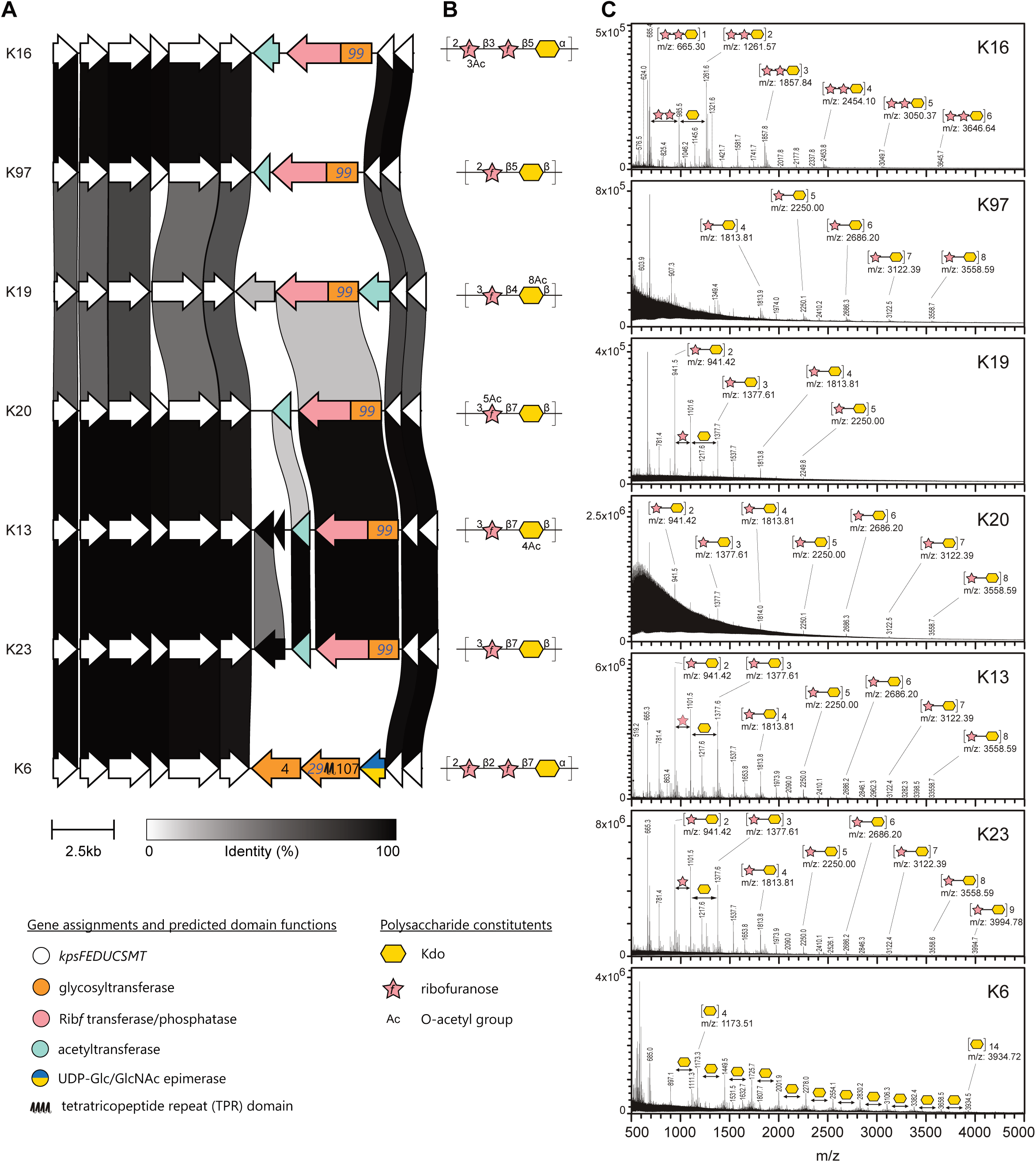
Mass spectrometry clarifies K antigen structures, related to Figure 2. (A) Capsule gene clusters from K serotype reference strains were compared and rendered with clinker, then colored according to gene assignments and predicted domain functions. Amino acid identity > 30% is displayed in greyscale between opposing genes. (B) Published polysaccharide repeat unit structures for each K serotype. (C) Deconvoluted MALDI-TOF-TOF spectra of partially hydrolysed and permethylated capsular polysaccharide (STAR Methods). Peak labels indicate the assigned structure and corresponding expected m/z species calculated in GlycoWorkbench 2. We note that O-acetyl groups are expected to be lost during the permethylation procedure and were not considered when assigning peaks.

**Figure S3.**
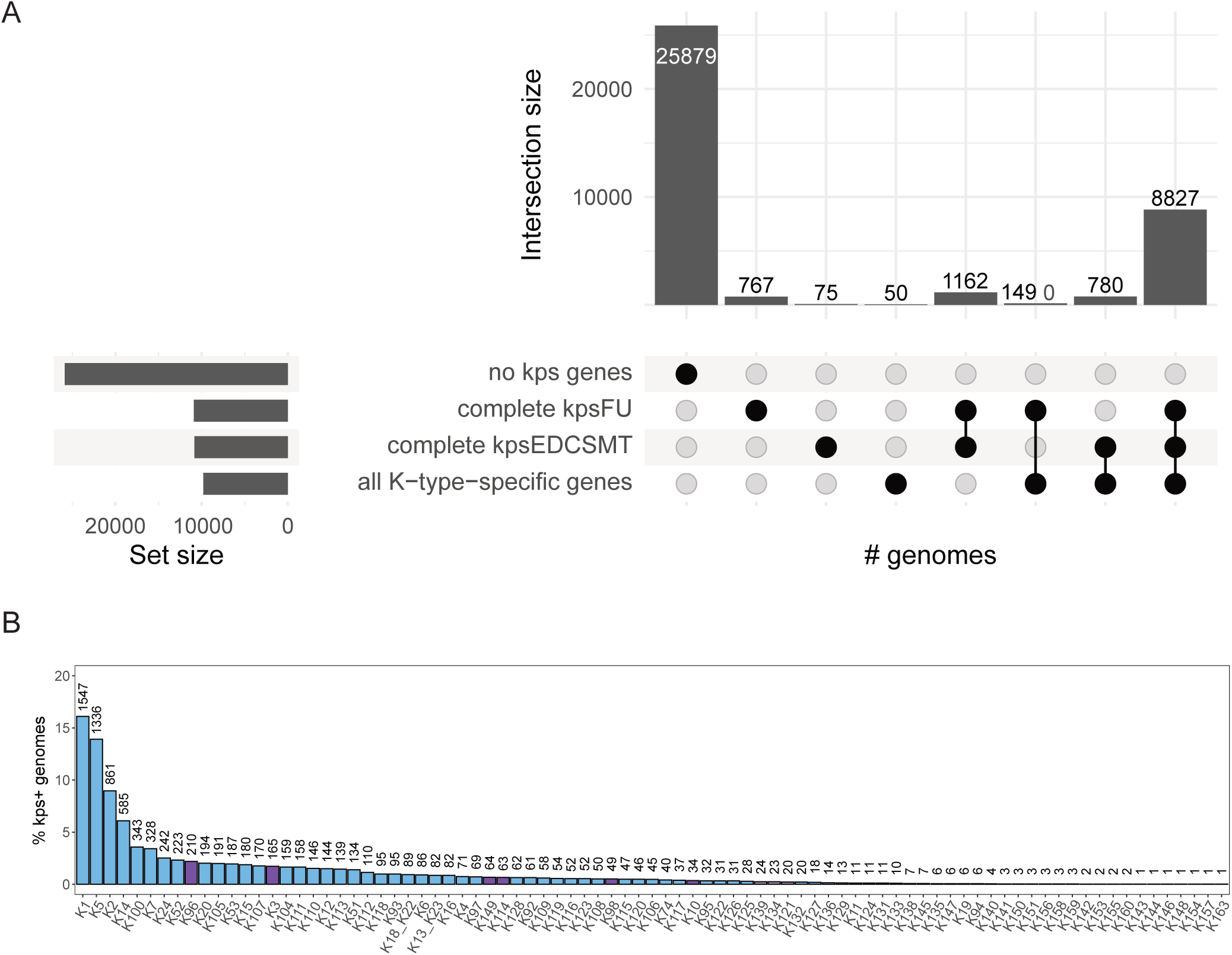
Evaluation of kTYPr on the RefSeq source data set. (A) Upset plot of kTYPr results on 37,723 *E. coli* genomes from RefSeq, showing the distribution of *E. coli* genomes with complete and incomplete capsule biosynthesis loci. (B) Absolute counts of K-types identified in the RefSeq data set.

**Figure S4.**
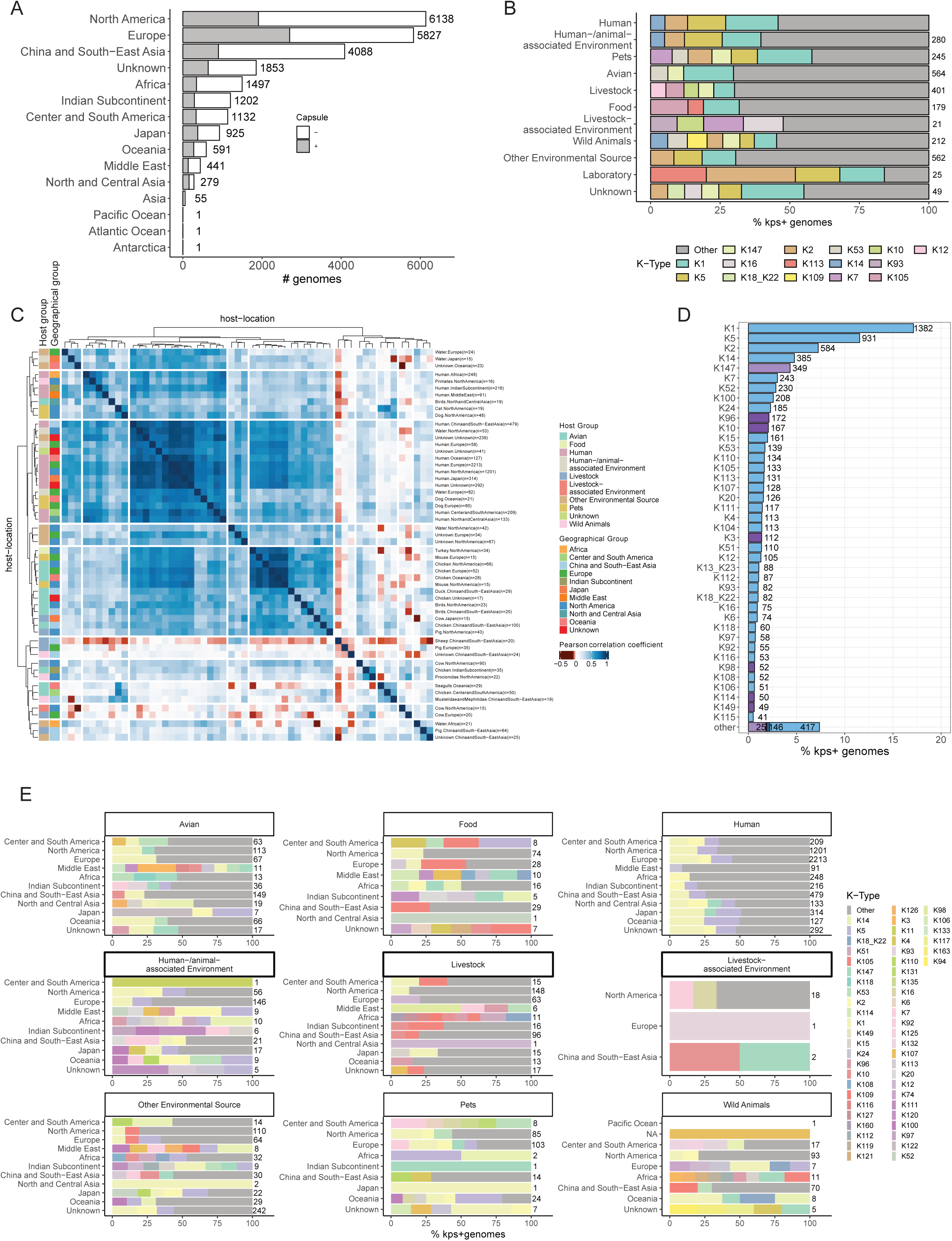
Global diversity of K-types, related to. Figure 3. (A) Fraction of capsule*-*positive genomes (defined as containing a complete *kps* locus, STAR Methods) across geography. (B) K-type proportion in hosts and environments. K-types with frequency < 5% are annotated as “other”. (C) K-type proportion in individual hosts and environments. K-types and sample origin are clustered according to Pearson’s correlation. Host-environment combinations with < 15 occurrences were not considered. K-type groups are color-coded as in Fig. 3C. Hosts are color-coded according to their lower resolution groups shown in Fig. 3B. (D) Proportion and absolute counts of K-types, color-coded according to groups as in Fig. 3C. K-types with proportion < 0.5% were annotated as “other”. Only *kps*-positive genomes were considered. (E) K-type proportion across geographic regions, grouped by hosts and environments. K-types with frequency < 8% are annotated as “other”. Only *kps*-positive genomes were considered.

**Figure S5.**
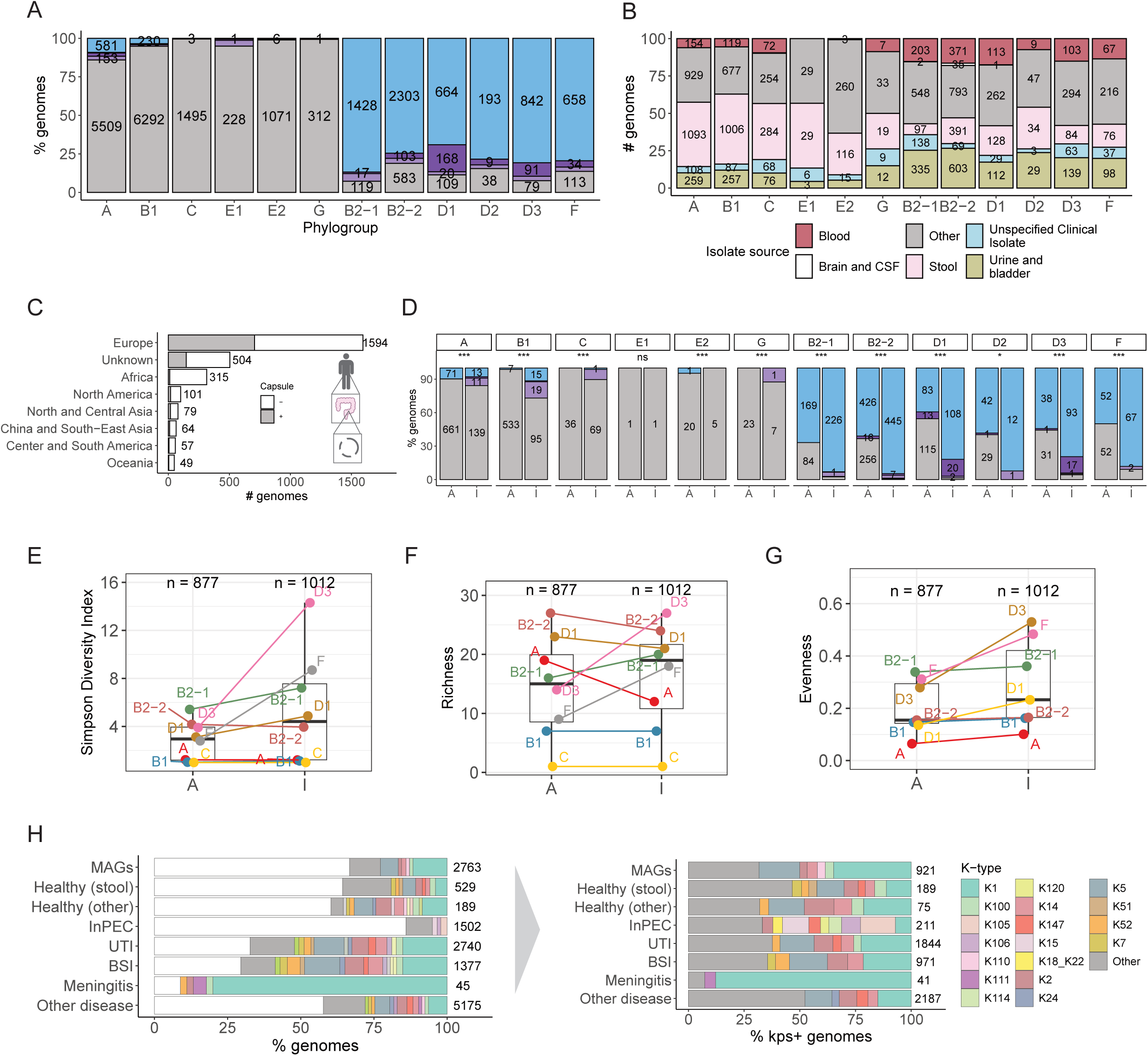
K-type associations with phylogroups and human health and disease, related to Figure 4. (A) Proportion of capsule groups across phylogroups in *E. coli* genomes from all hosts and environments. Groups are color-coded as in Fig. 3C. (B) *E. coli* human isolate source in the collection, grouped by phylogroup (n=11,537). 20 genomes to which no phylogroup could be assigned were removed. (C) Fraction of capsule*-*positive genomes (defined as containing a complete *kps* locus, STAR Methods) in *E. coli* human gut metagenomes. (D) Capsule group proportion in each phylogroup in asymptomatic carriage (A, corresponding to 2,762 *E. coli* MAGs from healthy individuals) and invasive *E. coli*-associated disease (I, 1,178 genomes of isolates from blood or cerebrospinal fluid from NCBI and 259 from blood from a published study on urosepsis), STAR Methods. ns p > 0.05; * p ≤ 0.05; ** p ≤ 0.01; *** p ≤ 0.001 (Fisher’s exact test). (E-G) K-type diversity (expressed as Simpson index, E), richness (F) and evenness (G) of phylogroups. The same data as in D) were considered, filtering out phylogroups with ≤ 25 genomes in each group (asymptomatic, A or invasive, I). The number of capsule-positive genomes in the two groups is indicated above each boxplot. Box limits correspond to first and third quartiles, with the median marked, and whiskers extending to the most extreme data points up to 1.5 times the interquartile range (IQR). (H) K-type proportion across health groups defined as in Fig. 4B, considering all (left) or only capsule-positive genomes (right). K-types with proportion < 1% (left) or < 3% (right) are grouped as “other”.

**Figure S6.**
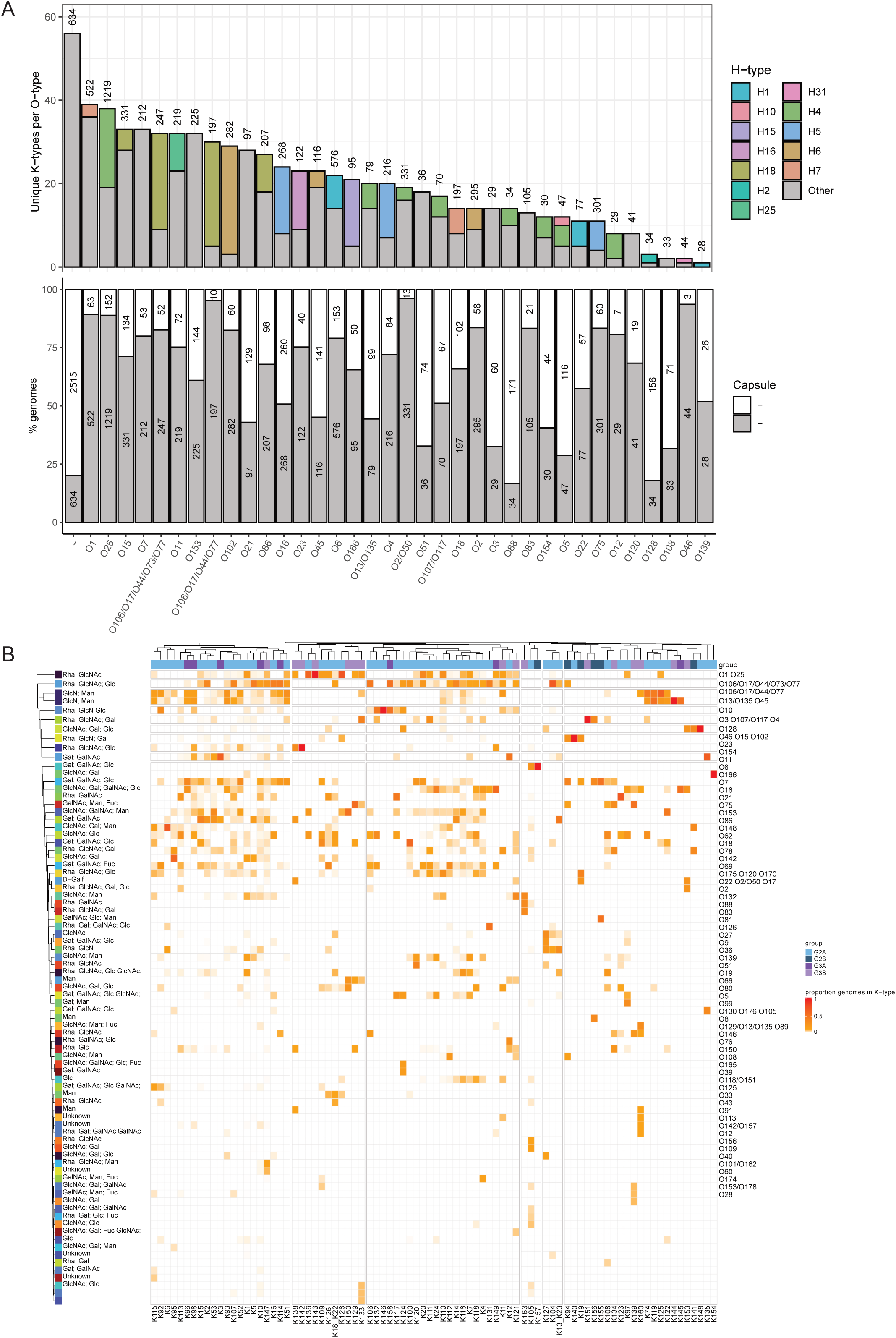
K-type associations with O and H antigens. (A) Proportion of *kps*-positive genomes (top) and the number of unique K-types (bottom) per O-type. H-type composition is indicated for each O-type (bottom). Only O-types with > 25 genomes are shown. H-types with proportion < 30% are grouped as “Other”. (B) O-type proportion for each K-type. K- and O-negative cases are not shown. Rare O-types (total proportion across all K-types ≤ 0.032, as first quartile of distribution) are not shown. K-type groups are color-coded as in Fig. 3C. O-type sugar composition was obtained from ECODAB.^2,72^ Only unique sugars are listed. K12 co-occurrence with rhamnose-containing O-types is highlighted.

**Figure S7.**
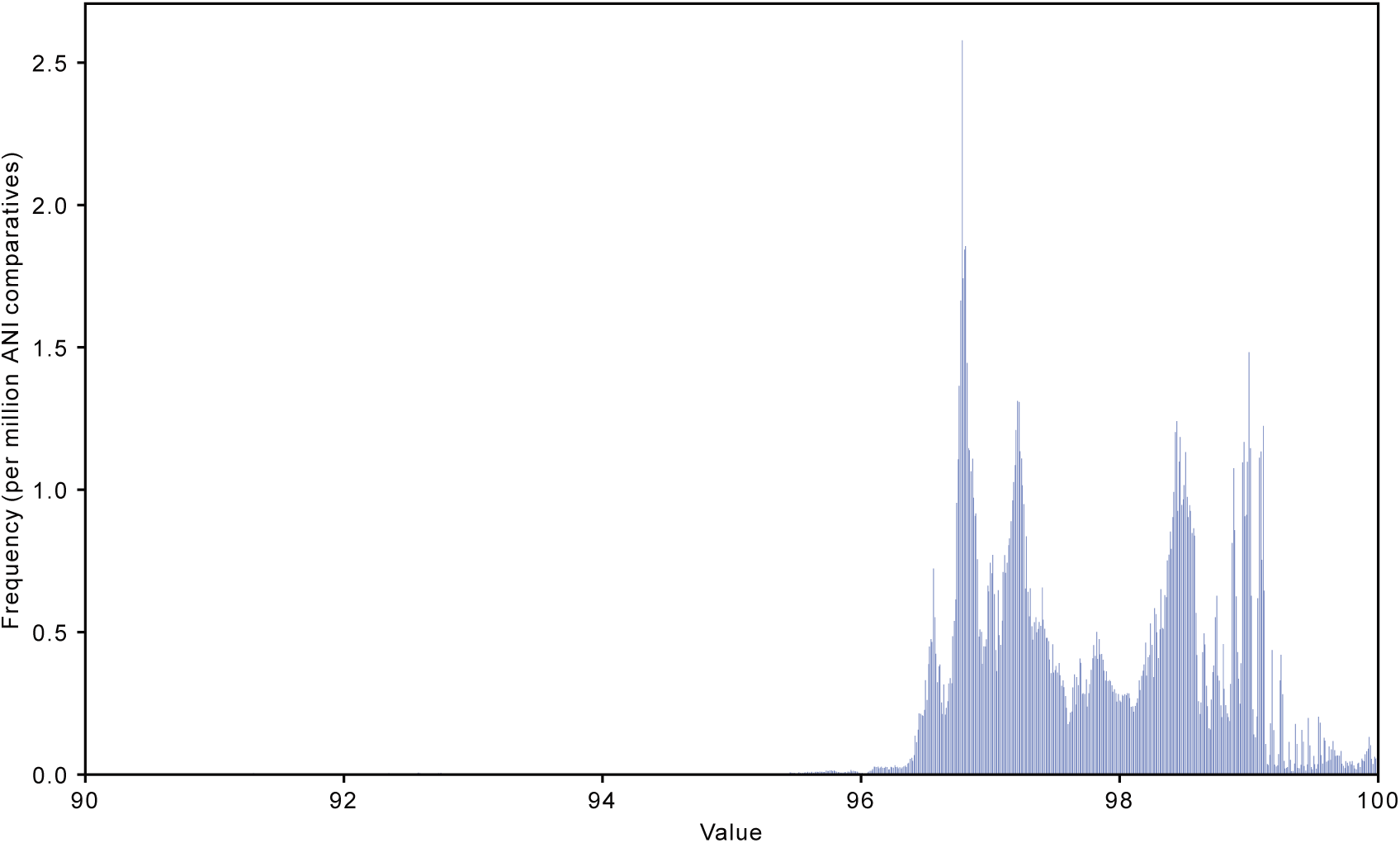
Distribution of ANI scores across genome comparisons. For 778,526,070 all-vs-all ANI comparisons, we show the frequency distribution (normalized per million pairs) using 1,500 bins for all the ANI values (X-axis). The 98% threshold was selected based on the separation of the two main modes of the distribution.

## STAR Methods

### Sequencing and genome assembly for K antigen reference strains

Strain acquisition, growth, DNA extraction, sequencing, and assembly of genomes is described (Roese Mores et al., Genome Announcements, 2025).

### Developing a catalog of capsule gene clusters

To survey the diversity of ABC transporter-dependent capsules in *E. coli*, we cataloged the *kps* gene clusters in a collection of 37,732 *E. coli* genomes available from RefSeq (downloaded 19^th^ May 2024). We used *kpsC* as a marker gene due to its large size and central role in capsule biosynthesis.^7,14,17^ *kpsC* positive genomes were identified with BLASTN (version 2.16.0)^73,74^ with the parameters `-max-target-seqs 100000 -max_hsps 1`, using *kpsC* genes from K1, K3, K11, K15, and K19 reference clusters as queries. We extracted the *kpsC* flanking region +/-30 kb from 11,834 genomes. 130 sequences where *kpsC* occurred on a contig of <8 kb were excluded. The remaining sequences were dereplicated at 90% identity and coverage using `mmseqs cluster` algorithm (MMseqs2 Version 15-6f452)^75^ with the parameters `–min-seq-id 0.98 -cov 0.9 cov-mode 0 -s 7.0`. The unique representatives (n=2,822) were annotated using DFAST Web API^76^ with the parameter `dataset=ecoli` because of its sensitivity for detection of *kpsFEDUCSMT* genes. Filtering for the presence of all essential genes (*kpsEDCSMT*) yielded 1911 unique sequences with a potential *kps* locus. To account for K-specific genes often, but not always, being flanked by the common *kps* gene, we trimmed each sequence to an approximate cluster; including a minimum of 2 ORFs up- and downstream to the *kps* start and end coordinates, or 10 ORFs up- and downstream in cases where no candidate K-specific ORFs were identified within the bounds of *kpsFEDUCSMT*. Dereplication of the trimmed sequences using `mmseqs cluster` with `-min-seq-id 0.98` yielded 292 representative sequences. Each sequence was visually inspected to define the locus start and end coordinates, considering the organization of *kpsFEDUCSMT*, functional annotations (DFAST), and identification of common flanking genes. Dereplication of the manually trimmed sequences at 98% identity yielded a set of 190 unique *kps* locus sequences.

We further refined the capsule locus catalog based on gene content. To do this, we first generated a global protein catalog by clustering all serotype-specific ORFs (excluding transposase-related sequences) at 90% amino acid identity and coverage. Each resulting cluster was aligned using MUSCLE (v3.8.1511)^77^ and used to build a protein HMM (HMMER 3.4)^78^. In an iterative process, these HMMs were applied to automate identification of locus boundaries within each *kpsC* flanking region +/-30 kb. This approach ensured the inclusion of functionally related genes that may have been missed during initial manual locus curation, and it allowed us to merge loci with the same gene content. In later iterations, we transitioned to K-type specific protein HMMs, built exclusively with sequences assigned to that K-type by analysis with kTYPr. Exceptions to this rule are proteins with a common function across multiple K-types, i.e. RfbBDAC, NeuDBACE, and KpsFEDUCSMT. Because closely related gene clusters can produce serologically and chemically distinct polysaccharides – such as K1 and K92, which differ by a single ORF with 83% protein identity – we used a threshold of 90% identity across all ORFs for defining distinct K-types. This led us to a final set of 85 K-types defined by a set of 456 protein HMMs.

### Phylogenetic analysis of conserved *kps* genes

Phylogenetic relationships among the conserved *kpsEDCSTM* genes were inferred to support the classification of *kps* loci into distinct groups. Protein sequences corresponding to each conserved gene (*kpsD, kpsE, kpsC, kpsS, kpsT*, and *kpsM*) were individually aligned using MAFFT (v7.305)^79^ with default parameters. Phylogenetic trees were constructed using FastTree (v2.1.10)^80^ under the JTT+CAT model for maximum-likelihood inference. The resulting phylogenies were visualized using iTOL v7^81^.

### Definition and development of kTYPr

The kTYPr tool developed and utilized in this study is fully written in Python (v≥3.9) and can be downloaded and installed following the details in https://github.com/SushiLab/kTYPr. This bioinformatic tool accepts two types of input in FASTA format: genomes or user-provided translated gene sequences. If a genome is provided, whether a complete genome or a metagenome-assembled genome, the analysis begins with gene prediction using *Pyrodigal* (v3.4.1) and its find_genes function^82,83^. Once a set of predicted protein sequences is obtained, kTYPr employs *PyHMMER* (v0.10.11) ^84,85^ to scan for the presence of *kpsC* within the sequences. If a *kpsC* hit is detected, the tool extracts a ±30 kb region flanking the start and end coordinates of the hit. At this stage, no thresholds or best-hit filtering are applied, meaning that if multiple *kpsC* genes are present, regardless of score, all identified *kpsC* candidates (if multiple) and flanking genes are retained for further analysis.

The extracted subset of genes is profiled against our custom catalog of 456 HMMs, each linked to a known K-type, generating an initial kTYPr output table. This table, called ‘all hits’, includes all identified genes, their corresponding HMM hits, and key metrics such as bit scores. HMM hits are then filtered according to custom bit score thresholds. By default, thresholds are set to 60% of the maximum bit score observed for each HMM in the RefSeq collection. In a handful of cases the thresholds were decreased to accommodate highly polymorphic genes where the observed variation is known (e.g. the K5 putative sulfotransferase, *kfiB*^86^), or not expected to be critical for K antigen production (e.g. variation in length of the coiled-coil regions in the putative methyltransferase genes present in many group 3 clusters). On the other hand, the thresholds for dTDP-rhamnose biosynthesis genes (*rfbABCD*) were increased to limit false positive hits to homologs present in many *E. coli* O antigen biosynthesis gene clusters. After applying these thresholds, the filtered set of hits is analyzed to determine the presence or absence of the conserved region, based on identification of *kpsEDCSMT* and *kpsFU* genes (assigned a value of 1 if all genes are present, 0 otherwise). The same strategy is applied to each K-specific gene set associated with different K-types. The final output table reports the presence of the conserved region and, for each K-type, whether the cluster is complete, along with its accumulated bit score.

Additionally, a separate set of columns designates the best K-type, prioritizing first the completeness of the cluster and then the highest accumulated bit score. This approach allows users to explore multiple or nearly complete clusters while still identifying the most probable gene-based K-type. The outputs are a detailed kTYPr report, genbank file for the extracted cluster, and an HTML file with comparison of the query cluster to the predicted or closest match K-type cluster.

Alternatively, to enable kTYPr to run on incomplete genomes, we have implemented a ‘whole-genome’ mode as an alternative to the ‘flanking mode’ described above. In this mode, the initial step of selecting neighboring genes around *kpsC* is skipped, and instead, every gene in the genome is evaluated throughout the process. Furthermore, the described bioinformatics process operates at the single-genome level; however, kTYPr also supports a batch mode (see https://github.com/SushiLab/kTYPr). This mode allows users to provide a list of genome file paths, enabling the same analytical steps to be applied across an entire genome collection. Additionally, multiprocessing capabilities are available to enhance performance, ensuring efficient processing of large datasets. In this mode, the final output consists of a comprehensive table that consolidates predictions for the entire collection, streamlining downstream comparative analyses.

All analyses presented in this study were performed using kTYPr in ‘flanking mode’, considering the custom definition of curated hit cutoffs, and in batch mode.

### Definition of reference gene clusters

For each of the 85 defined K-types, we assigned a reference gene cluster. These were assigned first based on kTYPr results, using the cluster with the highest accumulated bit score for each K-type. The assigned reference was controlled by clinker comparison with all clusters (or a representative set of all clusters identified using MMSeqs2 with a minimum sequence identity threshold of 90%) to ensure that the gene organization was typical of the majority of clusters assigned to the given K-type. We note that the reference gene cluster is never taken from the corresponding K antigen reference strain, though in some cases they may be identical.

The reference gene clusters were compared in clinker, and K-specific ORFs extracted from the reference gene clusters were used for functional annotation of the gene clusters (**Fig. 2**).

### Functional annotation of ORFs

Functional annotation of serotype-specific ORFs was performed using InterProScan (v5.62-94.0)^87^ and dbCAN3^42^ with default parameters. InterProScan result columns ‘orf_id’, ‘length’, ‘domain_start’, ‘domain_end’, ‘DB’, ‘DB_hit’, ‘DB_hit_description’, ‘e_value’, ‘gene_ontology’ were taken to group results by ORF identifier while preserving the information by the database were the hit was retrieved. This tab-delimited output and the standard output from EggNOG were then combined by ORF identifier for further comparative and analysis.

Protein structure predictions were performed using ColabFold (v1.3.0).^88^ The AlphaFold2 version used within ColabFold was v2.1.14.^46^ Multiple sequence alignments (MSAs) were generated using MMseqs2.^89^ To maximize sequence coverage, searches were conducted against the UniRef30^90^ and ColabFold environmental^91^ databases. MSAs were obtained using the colabfold_search command with default parameters, specifying both databases for alignment.

Protein structure predictions were then performed using the precomputed MSAs with the colabfold_batch function. Five AlphaFold2 models were generated per query, with a maximum of three recycles. Model confidence was ranked based on predicted local distance difference test (pLDDT) scores, and structures were retained according to this ranking. Template-based modeling and Amber refinement were not used. The models were generated using an ensemble size of one, and the pairing mode was set to “unpaired+paired” to optimize MSA depth. The MMseqs2 search mode included both UniRef30 and environmental databases to improve structural predictions.

Serotype-specific ORFs were clustered according to sequence and structural similarity using MMseqs2^75^ and Foldseek^47^, respectively. Sequence-based clustering was conducted using MMseqs2 (v14) with a minimum sequence identity threshold of 30% and 90% coverage. Clusters were generated using single-linkage clustering, ensuring that closely related sequences were grouped while maintaining sensitivity to divergent homologs. Structural homology clustering was performed using Foldseek (v8), using a TM-score threshold of 0.6 and a minimum coverage of 90%. Additionally, ORFs were searched against the Protein Data Bank (PDB) ^92^ to identify potential structural homologs using Foldseek (v8) and default parameters.

### Purification, hydrolysis, permethylation and MS analysis of capsular polysaccharides

K antigen reference strains were cultivated to stationary phase in TB media^93^ at 37°C shaking at 180 rpm in Erlenmeyer flasks. Total surface polysaccharides, including lipopolysaccharide (LPS) and capsular polysaccharide (CPS), were extracted from approximately 25 g (wet weight) cell pellets according to the hot phenol/water procedure (Westphal and Jann, 1965)^94^ with slight modifications as follows. Membrane lipids were extracted from the biomass once with 500 mL of ethanol overnight with stirring, then twice with 500 mL of acetone for 2 h with stirring. The cell mass was recovered after each extraction by centrifugation at 2,500 g, 20°C, 30 min, with slow deceleration, supernatant was discarded. After extraction of membrane lipids, the cell mass was air dried and weighed, typically yielding 3-5 g dry weight. Dry cell mass was crushed to a fine powder with a spatula, then resuspended with 60 mL of 50 mM Tris-HCl pH 8.0 at 65°C under vigorous stirring in teflon coated centrifuge buckets. 60 mL of preheated phenol at 65°C was added to the bucket, vigorous stirring of the aqueous-phenol mix continued for 15 min at 65°C. The aqueous phenol mix was then cooled in an ice bath for 10 minutes, then centrifuged for 45 min at 4°C with slow deceleration to reduce disruption of the separated phases. The upper aqueous phase containing LPS and CPS was collected to a clean vessel. The remaining phenol phase was extracted a second time with 60 mL of 50 mM Tris-HCl pH 8.0 using the same procedure, and the upper aqueous phases were pooled. The pooled aqueous extract was dialysed 5 times over 5 days with 40-fold volume of water using 14 kDa Spectra/Por 4 regenerated cellulose membranes to remove phenol. The dialysed extract was lyophilized to dryness, weighed, and resuspended at 50 mg/ml in 20 mM Tris-HCl pH 8.0 with 1 mM MgCl_2_, and treated with 500 Units of Benzonase for 1 h at 37°C. Lipid A and short LPS molecules were removed by three successive extractions with Triton X-114 as follows. 2% by volume of triton X-114 was added to ice cold samples then vortexed until completely dissolved. After 10 minutes on ice, samples were warmed to 42°C in a water bath to trigger phase separation of triton X-114, then centrifuged for 10 min at 20,000 g in a hot rotor. The aqueous supernatant was removed to a fresh tube, taking care not to disturb the triton X-114 extracted material at the bottom of the tube. Capsular polysaccharides were then selectively precipitated by adding 1 volume of 10% cetryltrimethylammonium bromide (CTAB). The precipitated material was recovered by centrifugation for 15 min at 4,500 g, and dissolving the pellet in 7 mL of 1 M NaCl. Polysaccharides were then precipitated from 1 M NaCl by addition of 3 volumes (27 mL) of ice cold ethanol, and centrifugation at 4°C and 8,000 g for 15 min. Supernatant was discarded and the pellet was resuspended in 7 mL of water. Purified capsular polysaccharides were partially hydrolysed in 50 mM TFA using the following conditions:

| <u>K antigens</u> | <u>Hydrolysis conditions</u> |
| --- | --- |
| K6, K16 | 25°C overnight |
| K13, K20, K23, K97 | 40°C 1 h, then 25°C overnight |
| K19 | 60°C 1 h, then 25°C overnight |

Hydrolysates were neutralised by addition of 7 mL 50 mM NaOH, then adjusted to 50 mL total volume with 20 mM Tris-HCl pH 7.5. Capsular polysaccharide fragments were then loaded on a 10 mL Source 15Q column, washed with 20 mM Tris pH 7.5, and eluted in a step gradient up to 1 M NaCl. The eluted polysaccharides were dialysed against water, then lyophilized to dryness and resuspended in water.

Polysaccharide fragments were permethylated to enhance glycan ionization efficiency. For each sample, 50 µL (50 ug of polysaccharide) was dried in a Teflon-lined screw-capped glass tube. A slurry was prepared by grinding NaOH pellets in dry DMSO, and about 0.5 mL was then added to the sample, followed by about 0.5 mL methyl iodide. The mixture was shaken vigorously for 20 minutes at room temperature. The reaction was quenched by the dropwise addition of water, followed by extraction with 2 mL chloroform. The chloroform layer was washed repeatedly with water until clear, then dried under a stream of nitrogen. Permethylated glycans were dissolved in 20 µL of pure acetonitrile, and 1 µL was mixed with 1 µL of 2,5-dihydroxybenzoic acid matrix (10 mg/mL) on the MALDI-target plate. Samples were analyzed using a Bruker Rapiflex MALDI-TOF-TOF mass spectrometer in positive ion mode.

### Genome collection preprocessing and annotation

For the NCBI collection, we collected 39,460 genomes reported as *E. coli* from NCBI (October 11th, 2024) and removed all deprecated ones and duplicated accession numbers, leading to 32,043 genomes. O- and H-types were annotated using the tool ECTyper (v1.0) ^1^ and sequence types were determined using the MLST tool (v2.23) available at https://github.com/tseemann/mlst. The genomes were classified into phylogroups using a published Mash distance-based approach ^59^. For phylogroup C, an alternative reference genome was selected (GCF_001515725.1) from the Microreact database, based on its classification under multiple criteria and highest sequence scores ^56^. Each queried genome was assigned to the closest reference phylogroup using a Mash v2.3 distance cutoff of 0.04 ^95^. Metadata available from BV-BRC^96^ was further manually curated. Since NCBI genomes are biased towards hosts and environments (e.g. clinical isolates) and include time-series from the same source, we dereplicated all genomes according to a 98% ANI cutoff using the tool skani (v0.2.2) ^97^. The command skani triangle -l genome_list.txt --fast was used to generate an ANI percentage matrix by performing all-against-all genome comparisons within the collection (with genome file paths specified in the input text file). The fast mode was selected to enhance scalability, given that results for genomes with high N50 and >95% expected ANI, as in the case of this study, are comparable to those obtained using the standard mode ^97^. This matrix was then processed to define clusters of genomes with at least 98% ANI. This threshold was determined based on the distribution of ANI values across the matrix, which exhibited a bimodal pattern, with the 98th percentile marking the boundary between the two modes (Fig. S7). Coupling this information with metadata on sample origin, we considered as distinct genomes with distinct 98% ANI clustering,geographic origin, host type, gender, age and health state when available and unique in sequence type, O-, H- and K-type. This resulted in 22846 distinct entries which we used for all downstream, considering as K-positive genomes fulfilling the following criteria: (i) a complete set of conserved and variable genes in the *kps* locus; (ii) *kps* clusters present on the same contig, which led to removing K137, K152, K161, K162.

### Statistical inference on strain invasiveness

To assess K-type contribution to the invasive phenotype, the NCBI collection was filtered to include only human hosts and samples annotated as “Bloodstream Infection,” “Meningitis,” “Neonatal Bacteremia” and isolated from blood and cerebrospinal fluid (n=1118), or “Healthy” (n=773). This dataset was further integrated with 259 samples from bloodstream infections (Cuénod et al., 2023) and with publicly available, metadata-curated MAGs from the stool of healthy humans (n=2,763).^60^ A generalized linear model was fitted using the R package stats v4.5.0, with invasiveness encoded as binary outcome, and K-type, O-type, phylogroup, ST, host gender, age and geography as fixed effects. To prevent sparsity and improve model stability, categorical variables with fewer than 15 (for O-types) or 10 occurrences (for ST and H-types) were grouped as “Other”. To prevent complete separation, K-types with ≤ 5 occurrences in both groups (invasive and not) were grouped as “Other”. Fixed effect estimates were extracted and odds ratios (ORs) with 95% confidence intervals (CIs) were computed by exponentiating the model estimates. All analyses were conducted in R (v4.5.0).

